# CRISPRi-assisted *E. coli* strains increase success rate of burdensome construct cloning

**DOI:** 10.64898/2026.06.02.729553

**Authors:** Ian Faulkner, Cholpisit Kiattisewee, Brian Darst, Pansa Leejareon, Yasuo Yoshikuni, Jesse G. Zalatan, James M. Carothers

## Abstract

Genetic constructs meant for metabolic engineering in nonmodel microbes often use similar genetic parts to those familiar to *E. coli* work. The typical workflow is to clone these parts into plasmids in *E. coli* before they are transferred to the nonmodel host or its genome. In many cases, the metabolic burden of these constructs is stronger in the *E. coli* cloning phase of the workflow than in the eventual host, possibly resulting in mutation or other failure during cloning. Here, we apply generic knockdown of a range of popular expression systems, using CRISPR interference, by targeting guide RNAs to either promoters or RBSs that are commonly used in metabolic engineering. Generic targeting of a constitutive promoter series, combined with genome integration of CRISPR components, allows the use of only one or a few specific cloning strains to achieve strong knockdown of a wide range of constructs. Further, we present a recombinase-based workflow for easily adding guide RNAs with custom targets, so that users can knock down any desired promoter or ORF. Together, this group of strains comprises easy-to-use cloning strains meant for increasing success rates of difficult or burdensome cloning reactions, ultimately allowing more ambitious genetic constructs to reach their intended context.

## Introduction

As synthetic biology aims increasingly toward complicated genetic circuitry and challenging metabolism in nonmodel organisms, the genetic tools necessary for these aims can become more difficult to build. For example, engineers often initially build burdensome or toxic genes into plasmids for construction and propagation in *E. coli*, even if the constructs are intended for genomic integration or for use in nonmodel organisms. Instability, mutation, or other cloning failures often occur in this construction phase^1^, perhaps because the promoters in use express more strongly in *E. coli* than in the intended eventual host, or because the copy number of the cloning plasmid is higher than the intended genomic integration^2^. This is especially true if the expression is designed to be constitutive, as is often the case in metabolic engineering meant to eventually be scaled up^3^. Expression levels of heterologous genes in the eventual host can therefore be in effect limited by the *E. coli* cloning strain’s tolerance of the construct’s burden, even if that burden is artificially inflated in the cloning phase of construction.

The bottleneck of burden on the *E. coli* host during the cloning phase is not limited to work in nonmodel organisms or genomic integration cargos. Complex genetic circuits^4^ involving multiple logical motifs or separate modules^5^ might be large enough to require construction in multiple pieces. These pieces in isolation might impose more burden on the host cells than they would within the full circuit, for example if certain repressive elements or feedback inhibition are not yet in place. Additionally, plasmids meant for expression in a cell-free context^6^ might use very strong promoters and require medium or high copy number for concentrated preps. Finally, especially burdensome or toxic cargos, including some metabolic enzymes or expression system components like Cas proteins or T7 RNA polymerase, might require knockdown for efficient cloning even if their intended use is on a plasmid. And for these very burdensome cargos, even leaky expression from an inducible or activatable control system can be problematic during the cloning phase: in those cases, cloning success rate would benefit from diminishing this leak.

To allow higher expression of heterologous genes in their eventual context, a helpful strategy for the synthetic biologist is to minimize the burden experienced by the *E. coli* strain during the cloning phase, which increases the chance that the construct is assembled correctly and remains stable while it exists as a plasmid in *E. coli*. This strategy can take the form of stronger repression in inducible systems, nonnative polymerase/promoter pairs^7^, split expression systems^8^, direct knockdown of plasmid cargo^9^, and more. A particularly useful solution would be an *E. coli* strain with cargo knockdown that is agnostic to the specific promoter used and ORF contained within the cargo, so that one strain can be used off-the-shelf for diverse cloning applications.

Here, we present a platform for cloning diverse constructs in a small suite of E. coli strains, derived from NEB Turbo, that use CRISPR interference to knock down any construct using one of several standard genetic elements. These elements include a commonly used standard set of constitutive promoters (the Anderson series, BBa_J231xx, parts.igem.org), a commonly used ribosome binding site (Bujard)^10^, and a consensus σ70 promoter^6^, depending on user choice and application. Targeting these elements upstream of the ORF is understood to decrease CRISPRi efficacy relative to targeting the early part of the ORF itself^11^, but we consider promoter or RBS targeting to be a worthy tradeoff for the versatility of targeting an arbitrary number of constructs with just a small number of cloning strains. The result is a suite of CRISPRi-assisted Cloning Strains, also known as CCS or MAGIC (Moderate Assurance of Greatly Improved Cloning) strains, used in exactly the same workflow as a normal cloning strain but with drastically reduced construct burden.

Targeting the Anderson series of promoters is a particular strength of this approach, because sequence similarity across the series (despite the promoters’ wide range of strengths) allows any promoter in the set to be targeted by a single guide, with variable tolerance of mismatches. This is an advantage relative to previous approaches because it is more promoter-agnostic, allowing standard use of one or a few strains with a wide range of attempted plasmid construction. As part of reporting development of these strains, we present characterization of how well each strain knocks down each of these standard targets, to aid in the end user’s selection of which promoter to use in their construct and in which strain to clone it.

Finally, we present a platform for easy construction of new CCS strains that incorporate custom gRNAs for knockdown of custom targets. Because not all constructs use the relatively standard elements described above, gRNAs that target arbitrary sequences—such as other promoters, or any desired ORF—can be easily designed, synthesized, and cloned into an intermediate plasmid by the end user. The new gRNA can then be easily integrated into a modified CRAGE-Duet landing pad^12^, resulting in a custom-targeted CRISPRi-assisted cloning strain. We demonstrate this workflow and the capabilities of this approach to enable cloning that is otherwise difficult or impossible. Overall, such effective and versatile knockdown of genetic constructs undergoing the *E. coli* phase of construction should enable more effective and efficient cloning of even stronger and more complicated systems, ultimately improving expression level and therefore performance or production in the eventual host organism.

## Results

In the course of cloning circuit components^13^ in isolation, we noticed that certain strongly-expressing constructs could only be successfully cloned with enough of the circuit intact that knockdown of the output was in place. This became a viable enough strategy for cloning troublesome constructs that we began to build specific CRISPRi plasmids, to be included in cloning strains, to aid in other high-output contexts. Examples of such contexts include plasmids meant for expression in cell-free lysate or plasmids bound for nonmodel organisms in which promoters are less active, like *P. putida* (Figure 1). Initially, because of the helper plasmid from which the CRISPR components were expressed, the p15A origin and chloramphenicol resistance marker were off-limits for the cloning plasmid. Additionally, the helper plasmid was always co-purified with the cloned plasmid, complicating downstream processing.

**Figure 1.**
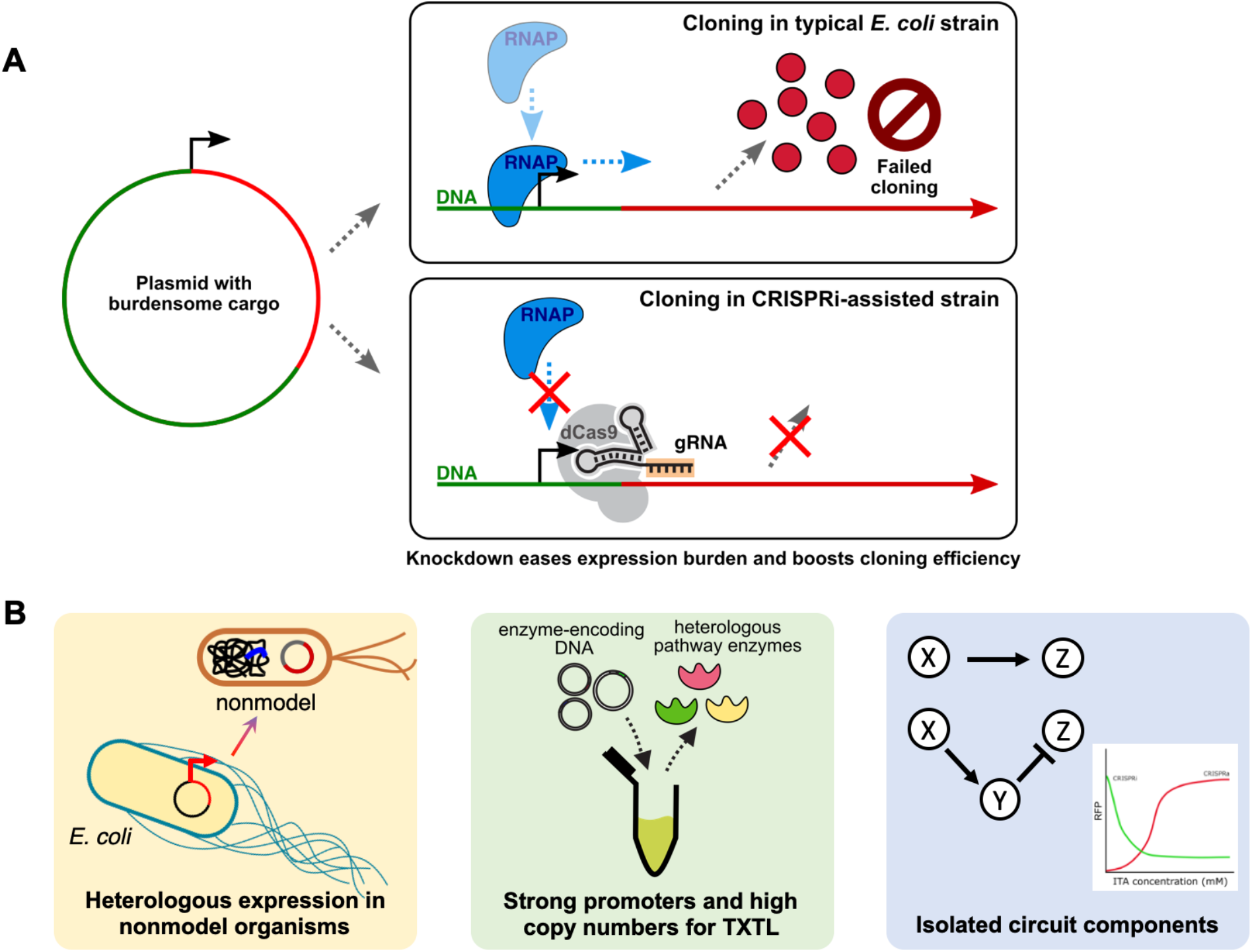
**A**. Cloning of burdensome plasmid cargos could fail in typical cloning strains because the cells are unable to overcome the toxicity or metabolic burden of any correctly-assembled plasmids, or because the selection pressure against correct plasmids reduces cloning efficiency. Knocking down expression of the plasmid cargo using CRISPRi can alleviate these issues and increase cloning efficiency. **B**. Such cloning is commonly difficult when the construct is intended for eventual use in nonmodel organisms, for integration into the genome, for cell-free expression, for genetic circuit construction, or for metabolic engineering.

To address the limitations of a plasmid-based approach, we then integrated the CRISPR components into the *arsB, rbsAR*, and/or *nfsA* sites around the genome (Figure 2) using λ-red recombineering, ensuring clonal purification of only the desired plasmid. While the Δ*arsB* site always contained dCas9 expressed by its endogenous *S. pyogenes* promoter (Supplementary Figure 1), the other two sites were initially devoted to expression of one or two single guide RNAs (gRNAs). The gRNAs were designed to target either the promoter or the ribosome binding site (RBS) that are commonly used in metabolic engineering and circuits work, as well as some combinations of the two. Targeting these control elements upstream of the gene’s ORF might not provide as strong of knockdown as an ORF-targeting gRNA, but it increases the versatility of a given gRNA for various cloning projects, since we tend to use similar promoters and RBSs across multiple applications. Also included in the strains were nontargeting control gRNAs and best-case knockdown control gRNAs targeting the open reading frame (ORF) of a measurable reporter, in this case mRFP1 or sfGFP.

**Figure 2.**
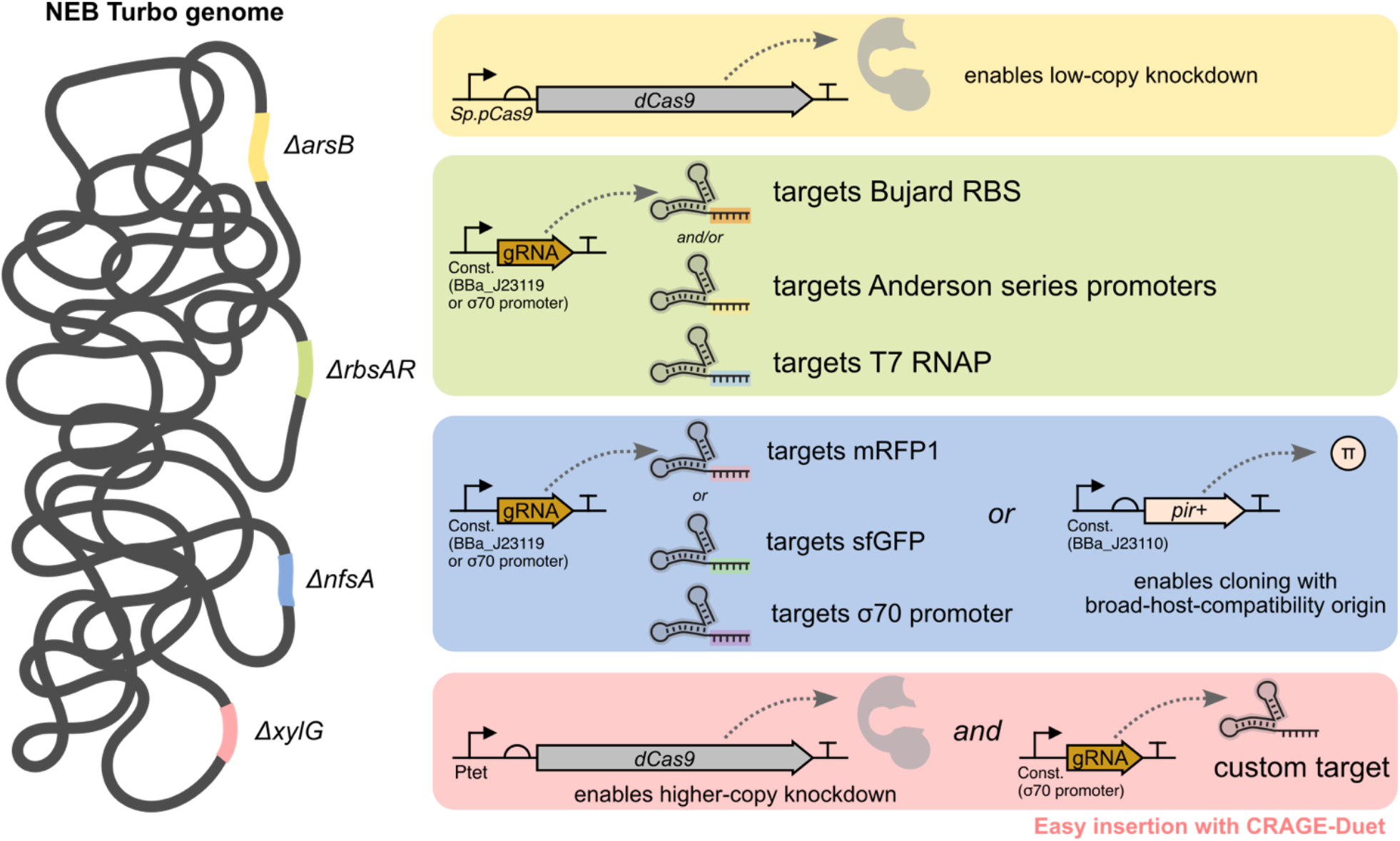
Starting with the NEB Turbo *E. coli* strain, dCas9 is always located in the site that disrupts *arsB*, under the control of the constitutive promoter from dCas9’s native context. One or two gRNAs are expressed from the Δ*rbsAR* site, under the control of a constitutive promoter that would not be self-targeting: for example, an Anderson promoter is not used to express an Anderson-targeting gRNA. One gRNA under similar constitutive control, or a gene expressing the pi protein, is located in the Δ*nfsA* site. Finally, CRAGE-Duet cargos (see Figure 5), usually including an extra dCas9 gene to supplement the Δ*arsB*-locus expression level and another gRNA meant to be easily designed to target a custom sequence, land in the Δ*xylG* locus.

Promoter-targeting gRNAs were designed to target either strand of the Anderson series of minimal promoters (BBa_J231xx, parts.igem.org) that cover a wide range of constitutive expression levels in many organisms. Because sequence-level differences between the different promoters in the series are minimal, the idea was that CRISPR complex binding to any promoter in the series would tolerate mismatches and still provide knockdown, even if the cloning strain contained a gRNA targeting only one of the Anderson sequences. This concept worked best using the target position on the top strand of the promoter (Figure 3A), because PAM-proximal positions contain very few mismatches across the set in that targeting window.

**Figure 3.**
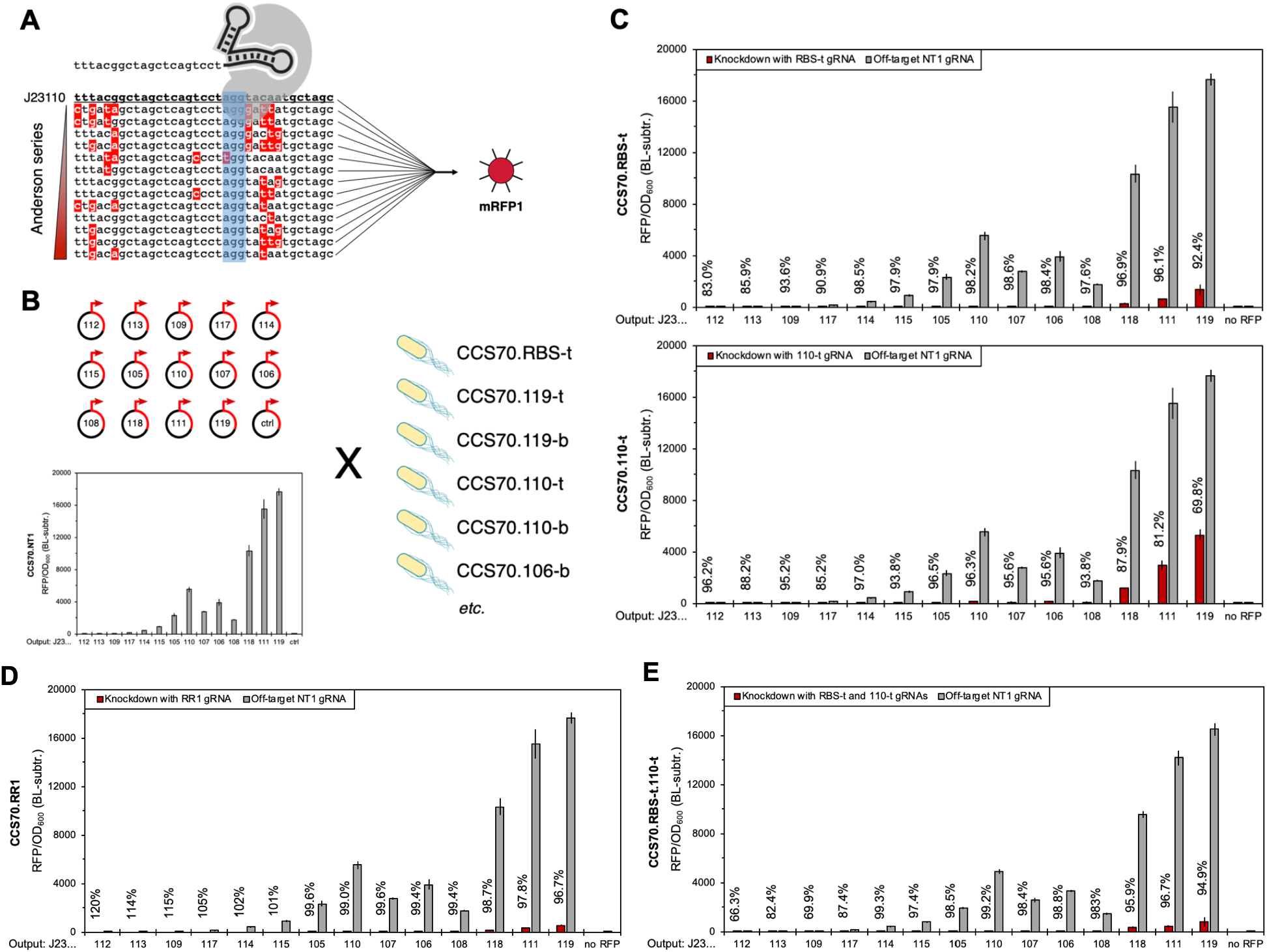
Guide RNAs targeting some Anderson promoters tend to knockdown any promoter in the series. **A**. When targeting using the highlighted PAM sequence, mismatches in the spacer sequence tend to be distributed toward the PAM-distal end. These positions are known to be less important for binding than PAM-proximal positions, which tend to be preserved across the set. This allows cross-talk from one gRNA across the series, therefore enabling one strain with integrated CRISPRi components to knockdown any promoter in the series to some degree. **B**. A set of low-copy plasmid reporters encompassing much of the promoter series driving RFP expression establishes each promoter’s baseline expression level before testing knockdown level in strains with various gRNAs integrated. Anderson promoter targets are available on the top strand (indicated by “-t”) and on the bottom strand (indicated by “-b”). **C**. Knockdown levels achieved by RBS-targeted and J23110-targeted gRNAs, each integrated into the genome in the strain indicated on the *y*-axes, across the series of reporters. Reporter expression has been knocked down to varying degrees depending on the DNA-binding mismatches tolerated by the CRISPRi complex in the CCS70.110-t case, but generally was reduced >80%. Knockdown is broadly stronger when using the RBS-t gRNA, but requires that the target plasmid use the Bujard RBS. These strains are shown as representative examples of mismatch characterization data; additional combinations are shown in Supplementary Figure XX. **D**. A control strain with the gRNA targeting the RFP ORF, though not useful for arbitrary cloning applications, provides a likely best-case knockdown level against which promoter-targeted and RBS-targeted knockdown should be compared. **E**. Combinations of gRNAs in the same strain achieve the higher of the two individual knockdown levels, but do not synergistically improve knockdown in this case (110-t combined with RBS-t). This dual-guide strain is therefore meant for increased targeting versatility rather than increased knockdown level. Grey bars indicate RFP expression from each promoter in the absence of knockdown due to an off-target gRNA, and red bars indicate repressed expression due to the gRNA(s) indicated in the strain name on the *y*-axis. Percentages indicate knockdown levels.

We built strains targeting only a few of the Anderson series members, relying on mismatch tolerance to allow knockdown across the whole set (Figure 3B). Strains targeting the output’s RBS were designed to target the Bujard RBS, raising concerns of widespread endogenous gene targeting due to sequence similarity. Growth in these strains appeared normal, however (Supplementary Figure 2), and there was no apparent loss in transformation efficiency (Supplementary Table 4).

Mismatch tolerance across a subset of Anderson promoters, the reporters of which each used Bujard RBS and the mRFP1 reporter, can be referenced in Figure 3C and Supplementary Figure 3. Also included are strains targeting the Bujard RBS, the consensus σ70 promoter (as another commonly-used constitutive promoter), the T7 RNA polymerase ORF, and the mRFP1 and sfGFP ORFs. In the Anderson series, the strain containing the 110-t gRNA generally performed best, but knockdown level depends somewhat on gRNA-promoter pairing. Users may consult the knockdown data in Figure 3C-E and Supplementary Figure 3 to determine the best strain to use for their construct. In general, the RBS-t gRNA targeting Bujard RBS provided strong knockdown in this series of reporters, and would also work well for constructs not using Anderson promoters. The S70a-t gRNA provided visibly strong knockdown of σ70-eGFP reporters in agar plate colonies (Supplementary Figure 5), but failed to quantitatively replicate that knockdown in liquid culture. Anecdotally, this strain does provide some benefit for cloning and transformation success, but concerns remain about plasmid stability in liquid culture.

To add compatibility with narrow-host-range origins, we then integrated either the pir+ or pir-116 gene into the nfsA site. Either gene allows maintenance, with variable copy number, of plasmids with the R6K origin through expression of the pi protein (Figure 4A). This compatibility is particularly important for cloning of genomic integration cargos, which often aim to use plasmids that are not replicable in the eventual host, and which often use strong promoters meant for genomic expression.

**Figure 4.**
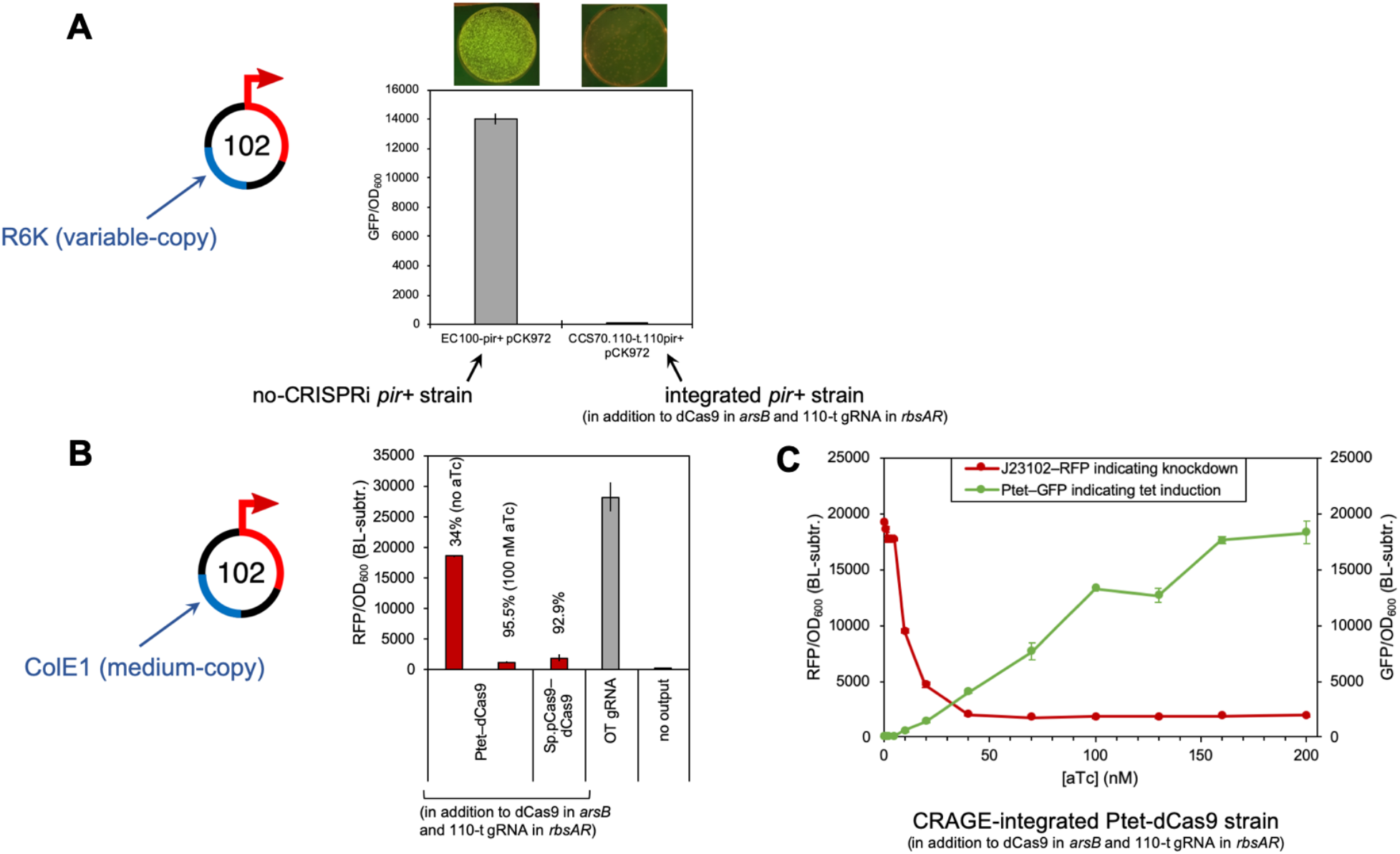
Expanding knockdown compatibility to R6K and medium-copy plasmid targets. **A**. Narrow-host-range plasmids, being often useful for genomic integration into hosts unable to replicate the plasmid, are an obvious use for knockdown cloning strains. The R6K origin of replication requires the pi protein supplied here by the *pir+* gene, integrated into the Δ*nfsA* site in the strain. This enables plasmid replication while the CRISPR components from the other genomic sites knock down an Anderson-promoter-driven GFP reporter. **B**. Medium-copy plasmids such as those using p15A (∼15 copies/cell) or colE1 (∼30 copies/cell) origins achieved poor knockdown in the strains with a single dCas9 locus (generally about 30%), but additional dCas9 expression from the Δ*xylG* locus boosts knockdown above 90%. Inducible dCas9 in this second locus provides the option to boost knockdown levels only when needed, as part of a general-purpose cloning strain. Grey bars indicate RFP expression from each promoter in the absence of knockdown due to an off-target gRNA, and red bars indicate repressed expression due to the 110-t gRNA. Percentages indicate knockdown levels. **C**. Titration of aTc inducer demonstrates the required level of extra dCas9 to achieve desired knockdown of a medium-copy RFP reporter. Maximal knockdown (red line, left *y*-axis) is achieved above roughly 50 nM aTc, which is not full expression from the P_tet_ promoter (green line, right *y*-axis).

Despite the high knockdown levels of low-copy plasmid targets, medium-copy plasmids such as those with p15A and colE1 origins could not be knocked down to a comparable degree by the initial strain (Figure 4B). This is presumably because the higher copy number presented too many targets for the pool of CRISPR complexes to adequately cover. After finding that additional expression of gRNA did not increase knockdown, we aimed to increase dCas9 levels by integrating a second locus of either P_S.py_–dCas9 or P_tet_–dCas9. To accomplish this additional integration in a new site, we adapted the CRAGE-Duet system^12^ to allow selection marker curing, then randomly integrated the landing pad into the CCS70.110-t strain (Figure 5A) using Tn5 transposase.

**Figure 6.**
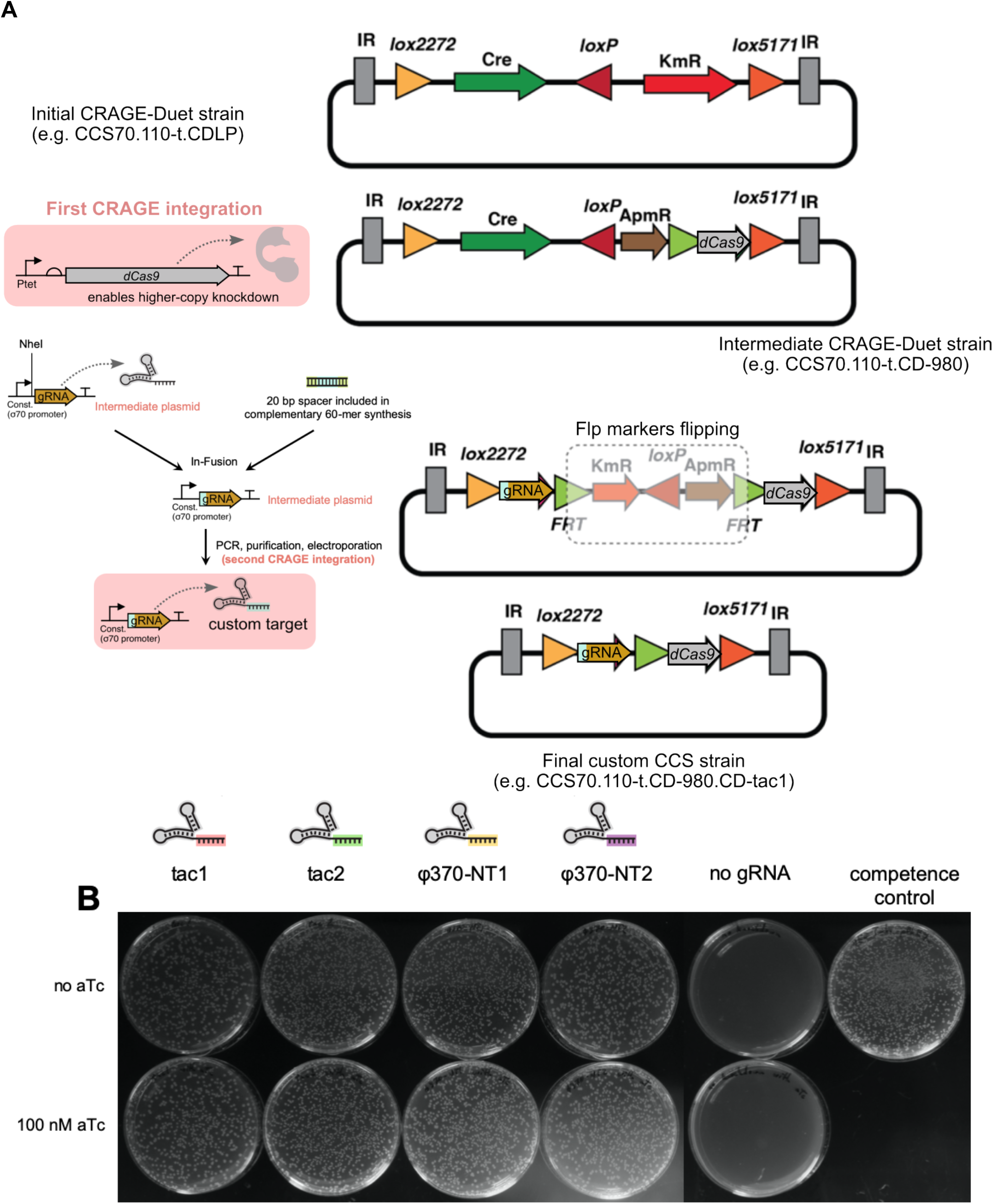
Curable CRAGE-Duet provides an easy route for integration of custom gRNAs. **A**. Once the landing pad is integrated, cells use the included *cre* recombinase to integrate electroporated cargo fragments into a first site and then a second. Here, the first site contains dCas9 and an apramycin resistance marker, replacing the original kanamycin resistance. The second site introduces another gRNA into the system as well as kanamycin resistance, replacing the recombinase. This gRNA can be easily designed as a custom target sequence by cloning a synthesized fragment into an intermediate plasmid. Naturally, curing the resistance markers from the genome is an important feature of a cloning strain, so the landing pad is arranged so that both apramycin and kanamycin resistance markers are situated between FRT sites for removal by FLP recombinase. **B**. Custom gRNA synthesis and integration allows for stronger knockdown (due to ORF targeting) of particularly burdensome constructs or of constructs that use different promoters or RBSs than the existing options. For example, the φ370 serine recombinase under P_tac_ control was difficult or impossible to clone without knockdown (second panel from right), but design and inclusion of one of several promoter- or ORF-targeting gRNAs (four left panels) enabled efficient cloning, with hundreds of colonies per reaction. Induction of extra dCas9 with aTc (bottom row) did not seem to improve efficiency, suggesting dCas9 was not limiting in this case. The rightmost panel depicts transformation of a control plasmid into the knockdown-free competent cells, to verify transformation efficiency.

Upon screening integrants for normal growth (about 6% of candidate colonies), the landing pad in the selected strain had randomly integrated into the xylG gene (Figure 2). The modified CRAGE landing pad allows integration of two cargos before curing kanamycin and apramycin resistance. While the original system also integrated two cargos, we accomplished subsequent curability by reorienting the resistance markers relative to the cargos and adding sites for FLP recombinase (Figure 5B or Supplementary Figure 6). Cargo addition is usually accomplished sequentially, so we designed the extra dCas9 locus to integrate into the first CRAGE site, leaving the second site for later integration of gRNAs. The most effective versions of this dCas9 integration were under the constitutive control of its native promoter or under the aTc-inducible control of the P_tet_ promoter (Figure 2), both of which boosted knockdown of a medium-copy target from about 30% to more than 90% (Figure 4B). For this reason, we recommend that the dual-dCas9 version of any CRISPRi-assisted cloning strain be the default version for routine use.

In order to replicate the CRAGE site in other strains with other knockdown targets, we found that λ-red recombineering with *xylG* homology arms was sufficient to move the entire landing pad between strains while preserving its genomic context. In this manner, we CRAGE-enabled CCS.RBS-t (for Bujard knockdown), CCS70.110-t.110-pir+ (for R6K compatibility), and CCS70.NT1 (for an off-target control strain) to provide additional targeting options in the dual-dCas9-loci versions of the cloning strains. We preferred to preserve the xylG locus in all of these strains because it would likely result in equal expression levels between strains in quantitative comparisons, and because each version of the strain would be pre-screened for normal growth. The result is a suite of strains that have the capability to induce additional dCas9 (and therefore strongly knockdown targets on medium-copy plasmids like colE1) and to receive custom-designed gRNAs as part of a simple workflow.

From any of the CRAGE-enabled strains, integration of any arbitrary gRNA is a simple transformation of a linear DNA template like a PCR product. Furthermore, custom gRNA spacer sequences for novel targets can be included in this linear fragment simply by synthesis of complementary 60-mer oligonucleotides followed by In-Fusion cloning into an intermediate plasmid (Figure 5A). We demonstrated this workflow by integrating one of several versions of custom gRNAs designed to knockdown the P_tac_ promoter or the ORF of a burdensome gene, the φ370 serine recombinase. After designing the spacer sequences, screening them for proper folding^14^, ordering the oligonucleotides and cloning them into a plasmid, and integrating the PCR product from that plasmid, we built a series of strains custom-designed for burden-reduced cloning of this particular gene. In the cases of all four guides tested, cloning success of the P_tac_–φ370 plasmid led to hundreds of colonies, compared to zero without the custom-targeted knockdown (Figure 5B) and in previous efforts.

In addition to the φ370 cloning example, we have used these strains to enable difficult cloning in many contexts across our lab. Some components of a feedforward circuit that were required to be expressed strongly could only be cloned with knockdown provided by the cloning strain or by the full circuit. A three-operon metabolic pathway for producing an aromatic product was planned to be integrated into the *P. putida* genome with strong promoters, but the cargo plasmids could not be cloned in *E. coli* without knockdown. Similarly, a dCas9-expressing construct meant for use in nonmodel purple non-sulfur bacteria required a strong-in-*E. coli* J23119 promoter, the cloning of which was enabled by a knockdown strain. The consensus σ70 promoter is also very strong in *E. coli*, and was often used in cell-free expression contexts but required initial cloning in a CRISPRi-assisted cloning strain. Across a wide range of applications, then, the relatively small suite of strains included in this work enables cloning that would be difficult or impossible without knockdown.

## Discussion

The strains presented in this work were built out of necessity due to the burden, promoter strength, and complication of constructs required for various projects. A major goal of the strain construction, then, was to provide a small number of options that can easily be used off-the-shelf for a wide range of applications. Given our common use of elements like Anderson promoters across multiple projects—combined with the mismatch tolerance of gRNA binding within that series—those promoters and some other common elements provided a great opportunity to build such widely-applicable knockdown strains. The high degree of knockdown still evident in the genome-integrated versions of the strains, relative to the original plasmid-based CRISPRi system, was a welcome development. Though we have largely surpassed the plasmid-based system, it still provides such strong knockdown and is so easy to build that it retains utility in certain systems, like recA-deficient strains or donor strains in conjugations. Its limitations remain, including copurification of the CRISPRi plasmid with the cloned plasmid, its use of chloramphenicol resistance, and some instances of CRISPRi plasmid instability—so most of our strain development focused on integrated versions.

The λ-red phase of strain development focused on previously characterized integration sites^15^, using sequential integrations starting with the most difficult: dCas9 in *arsB*. This strain, essentially NEB Turbo with dCas9 added, was the common starting point for various gRNA and *pir* gene integrations. Those variations generated the suite of cloning strains that knock down commonly used control elements. While the synthetic Anderson series of promoters is largely orthogonal, knocking down other features like the Bujard RBS and σ70 promoter carries a risk of crosstalk to native sequences across the *E. coli* genome, potentially putting cell health at risk. In our testing, however, detrimental effects in these strains were minimal: for example, CCS.RBS-t shows slightly lower endpoint density but similar growth rate and transformation efficiency compared to strains with other knockdown targets. And CCS.S70a-t does not appear to show effective knockdown in liquid culture, but is clearly effective in colonies on agar plates, which is perhaps more important immediately after transforming a cloned plasmid. Despite their potential targeting of native sequences, then, strains that target these RBS and promoter elements remain useful options.

Still, off-the-shelf options are not universally applicable to every desired construct, so the easy workflow for generating custom knockdown targeting was a welcome addition to the CRAGE-Duet phase of strain development. While originally intended to optimize dCas9 expression for various target copy numbers, the second integration site of CRAGE-Duet makes individual gRNAs or even gRNA arrays easy to add, and our addition of marker curability to the system keeps the resultant strains broadly applicable.

In some contexts, the common use of elements like Anderson promoters means that these strains are capable of knocking down some of their own components. For example, initial designs used J23119 to express the gRNA, resulting in negative feedback if that gRNA targeted any Anderson promoter. Interestingly, we found that these strains still seemed to work well despite these unintentional dynamic behaviors. Even so, we removed that possibility by changing Anderson-promoter-targeting gRNAs to be expressed by the σ70 promoter in the final strains. Similarly, the integration of the *pir+* gene driven by a J23110 promoter worked more reliably than the original promoter from commercial strains. But the CRISPRi cloning strains containing a 110-t gRNA are always knocking down expression of the pi protein from that gene, presumably affecting plasmid replicability. We observed no effect on the ability of those strains to host R6K plasmids, however, even while they were knocking down Anderson-promoter-driven plasmid cargo. Perhaps enough pi protein escapes knockdown either through leaky expression or initial expression before CRISPRi complex formation, such that R6K plasmids can still replicate at some copy number.

Another potential pitfall of promoter-targeted gRNAs in a CRISPRi system is that the spacer sequence in the gRNA expression cassette can itself act as a partial promoter. For instance, in the case of top-strand Anderson-targeted gRNAs in our system, the spacer sequence is exactly the -35 to -15 region of the nominally-targeted promoter. This could result in unintended transcription initiation 20bp downstream of the intended (and still active) TSS, resulting in turn in partial gRNA transcripts containing only a dCas9 handle. These handle-only RNAs would normally be worrisome for CRISPRi efficacy if the non-targeting handles were numerous enough within the RNA population, but in practice we have not seen diminished efficacy of strains that use top strand gRNAs. This was observed in terms of both knockdown efficacy (top-strand gRNAs tend to work better than bottom-strand gRNAs) and transformation efficiency (they work equally). In fact, the top-strand gRNA targeting J23110 (110-t) seems to be the strongest and most widely-applicable gRNA in the set.

Among all of the CRISPRi-assisted cloning strains built in this work, the most developed and widely applicable strains consist of a constitutive dCas9 gene, an additional inducible dCas9 gene, a pir+ gene, one or two of several gRNAs targeted to commonly used control elements, and a landing pad for easy addition of a custom-targeted gRNA if desired. We aim to make this suite of strains available to the community to enable engineering of more ambitious, more complicated, and more productive genetic systems especially in nonmodel bacteria. These diverse hosts are exciting chassis for engineering metabolism in the future, and many of them require further development of genetic toolkits before becoming sufficiently tractable. For example, the relatively well-understood nonmodel bacterium *P. putida* is a very suitable host for producing aromatic chemicals, but because the Anderson promoters that work both in *P. putida* and in *E. coli* tend to work more strongly in *E. coli*, and because of the copy number difference between *E. coli* plasmids and the *P. putida* genome, we could not clone a genome-bound multi-gene pathway for aromatics production without CRISPRi in the cloning strain. When we switched to a strain with knockdown in place, we effectively raised the ceiling of allowable promoter strength that we could clone, effectively increasing production from the eventual host *P. putida*. In this case, we relied on a standard set of promoters to control even a very burdensome genetic program, picking out the appropriate expression level for the eventual host organism and for genomic copy number—all while avoiding the roadblock of metabolic burden in the cloning phase. We hope the availability of these MAGIC CRISPRi cloning strains will enable similar development of genetic tools in more diverse and interesting microbes, and therefore engineered metabolism for real-world solutions to problems in bioproduction of chemicals and in other fields.

## Methods

### Plasmid cloning and sources of genes

Bacterial strains used in this study are described in Supplementary Table 1. Plasmids were cloned using standard molecular biology protocols and are described in Supplementary Table 2. Spacer sequences for gRNAs were screened for favorable folding parameters as previously reported^14^ and are described in Supplementary Table 3. S. pyogenes dCas9 was expressed using the endogenous Sp.pCas9 promoter. The *pir+* and *pir-116* genes were PCR-amplified from EC100D *pir+* (Biosearch, ECP09500) or EC100D *pir-116* (EC6P095H) cells.

### Strain construction using λ-red

Recombinase proteins were expressed at 30°C from pKD46^15^ using 0.23% arabinose at an OD_600_ of 0.8. After one hour of induction, cells were made electrocompetent, and a PCR product of the integration cargo flanked by homology arms was transformed. Outgrowth cultures were left at 37°C overnight before plating on LB agar plates with antibiotic selection. For moving the CRAGE-Duet landing pad between *E. coli* strains while preserving the *xylG* locus, the landing pad was made into a linear fragment with homology arms by colony PCR using forward primer gtaattaaagacggattccacaaagagagc and reverse primer cgtcacaatgatggtaagtggcaaag. After verification of integration, the marker was cured with pCP20 transformed by heat shock, the plasmid was cured by growth at 37°C, and the final strain was screened for antibiotic sensitivity.

### Strain construction using CRAGE-Duet

The landing pad for CRAGE-Duet, derived from pW37^12^, was linearized and randomly integrated using the EZ-Tn5 Transposase kit (Biosearch, TNP92110). Integrants were picked as single colonies from LB agar into liquid LB, and growth rate was assessed compared to wild-type using a kinetic growth assay on a Biotek Synergy HTX plate reader at 37°C (see culture conditions section, below). CRAGE-Duet cargos for integration into the landing pad were electroporated as whole plasmids (if the recipient strain did not contain a *pir+* or *pir-116* gene) or as purified PCR products encompassing the cargo and the flanking *lox* sites (if the recipient strain contained a *pir+* or *pir-116* gene). Outgrowth cultures were left at 37°C overnight before plating on LB agar plates with antibiotic selection. After sequential integration into both sites of the landing pad, markers were cured with pCP20 transformed by heat shock, the plasmid was cured by growth at 37°C, and the final strains were screened for antibiotic sensitivity.

### Custom gRNA workflow

Arbitrary sequences can be targeted using tools like Benchling for gRNA design; alternatively, known NGG PAMs can be used to determine the immediately preceding 20 bp of sequence on the same strand as the spacer sequence for the new gRNA. Then, the In-Fusion insert for the intermediate plasmid cloning can be ordered as complementary oligos from a DNA synthesis company like IDT with the following sequences flanking the new spacer: GATAATGGTTGCAGCTAGCxxxxxxxxxxxxxxxxxxxxGTTTTAGAGCTAGAAATAGC. This insert should be mixed, diluted about 1:1000, and melt-annealed before the reaction, and the backbone is pIDF115 digested with NheI. After successful In-Fusion, transformation, and screening of clones, the insert for CRAGE-Duet integration can be amplified by PCR (even colony PCR to save time) with forward primer ggcgcgcccagctgtctagg and reverse primer atttaaatcgtaattattggggacccctgg. This insert can be gel-purified and electroporated directly into the recipient strain for integration. We performed this workflow for tac1, tac2, φ370-NT1, and φ370-NT2 gRNAs (Supplementary Table 3).

### Culture conditions

Cloning of test plasmids was performed using restriction-ligation or In-Fusion, and 5 uL of each reaction was transformed using heat shock. For already constructed and miniprepped plasmids, 1 ng was used for each heat shock transformation, with concentration before dilution measured on a Nanodrop. Outgrowth cultures were kept at 37°C for 90 minutes before plating on LB agar with antibiotic selection. Single colonies were picked into triplicate cultures of 400 uL LB with appropriate antibiotics in a 2 mL 96-deepwell plate (Axygen) and shaken overnight at 37°C on an orbital microplate shaker (Heidolph). Endpoint cultures were measured in a Biotek Synergy HTX plate reader with 200 uL in each well of a flat, clear-bottomed black plate, measuring OD_600_, GFP fluorescence (excitation 485 nm, emission 528 nm), and RFP fluorescence (excitation 540 nm, emission 600 nm). These overnight-grown cultures were also diluted 1:100 into 200 uL of fresh LB with appropriate antibiotics in a flat, clear-bottomed black plate for a kinetic growth assay, in which the plate was incubated with shaking at 37°C for 16 hours with OD_600_, GFP, and RFP measured every 5 minutes.

