## Supplemental Figures for "CRISPRi-assisted *E. coli* strains increase success rate of burdensome construct cloning"

for

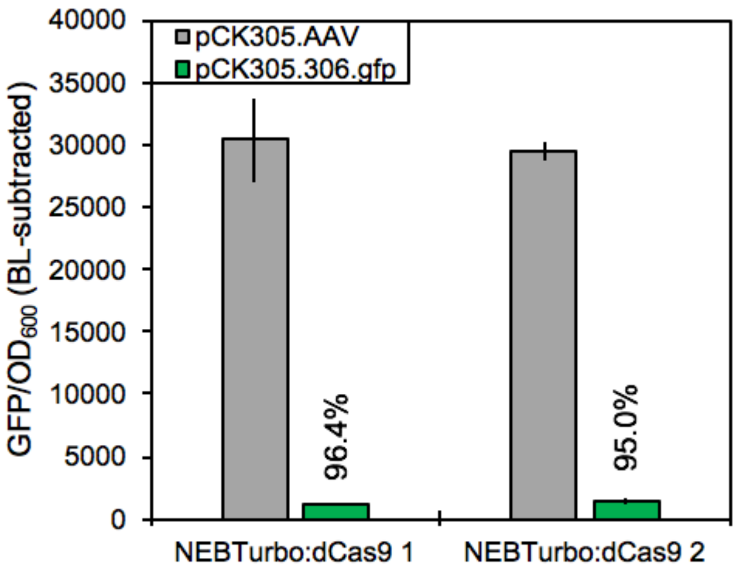


Supplementary Figure 1. Functional dCas9 integration into NEB Turbo. In this case the dCas9 is driven by the native Sp.pCas9 promoter and the gRNA is expressed from a plasmid using the J23119 promoter. The grey bars express an off-target gRNA, while the green bars express a GFP-targeted gRNA. Percentages indicate knockdown levels.

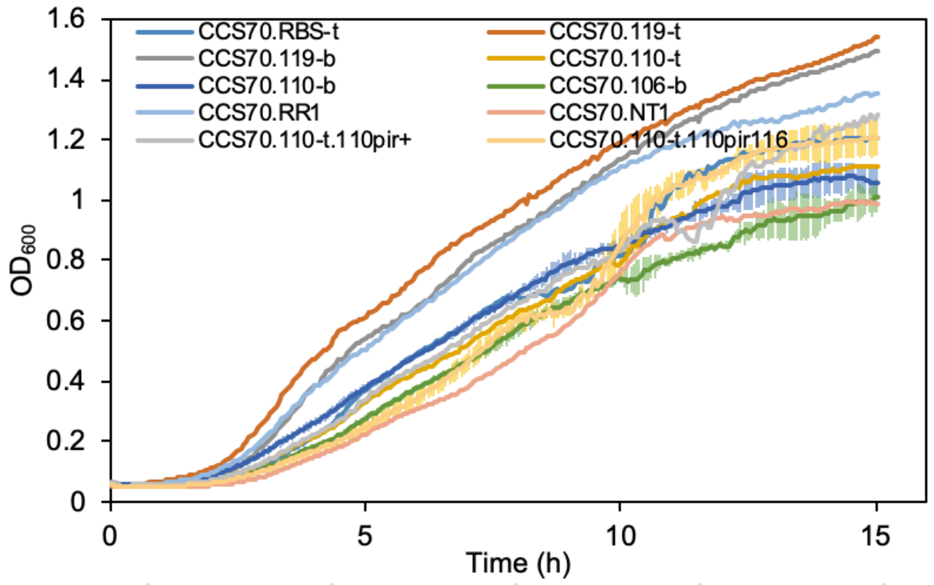


Supplementary Figure 2. Growth curves of CCS strains targeted to Anderson promoters or the Bujard RBS. The strain, and therefore the gRNA target, is indicated on the *y*-axes. The variously dark red lines indicate plasmids hosted by those strains with a wide range of promoter strengths within the Anderson series, and are best burden-matched to the “no RFP” comparison, an empty plasmid (black line).

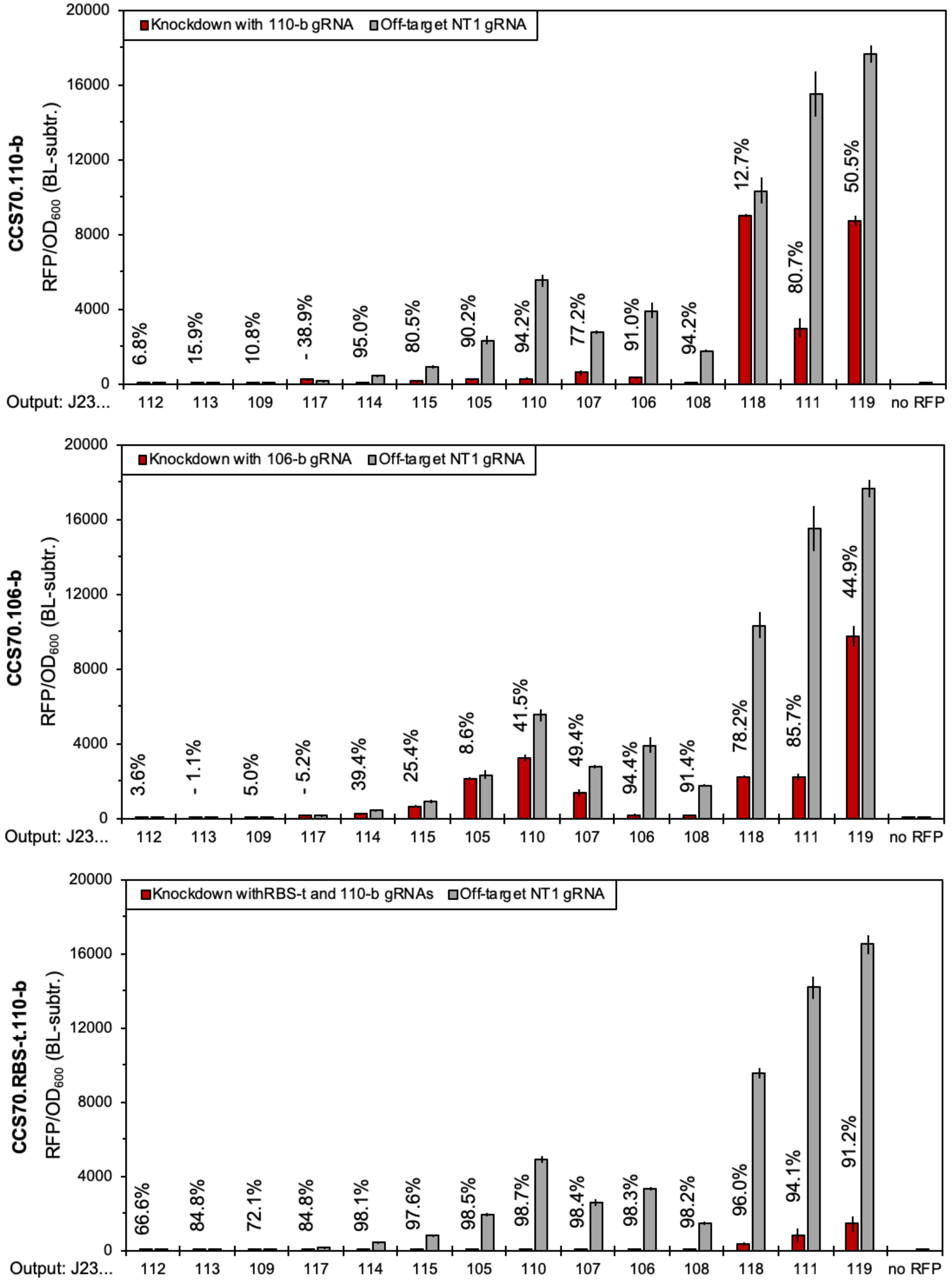

Knockdown with 110-b gRNA
Off-target NT1 gRNA
CCS70.110-b
RFP/OD600 (BL-subtr.)
20000
16000
12000
8000
4000
0
6.8%
15.9%
10.8%
- 38.9%
95.0%
80.5%
90.2%
94.2%
77.2%
91.0%
94.2%
12.7%
80.7%
50.5%
Output: J23...
112
113
109
117
114
115
105
110
107
106
108
118
111
119
no RFP
Knockdown with 106-b gRNA
Off-target NT1 gRNA
CCS70.106-b
RFP/OD600 (BL-subtr.)
3.6%
- 1.1%
5.0%
- 5.2%
39.4%
25.4%
8.6%
41.5%
49.4%
94.4%
91.4%
78.2%
85.7%
44.9%
Knockdown withRBS-t and 110-b gRNAs
Off-target NT1 gRNA
CCS70.RBS-t.110-b
RFP/OD600 (BL-subtr.)
66.6%
84.8%
72.1%
84.8%
98.1%
97.6%
98.5%
98.7%
98.4%
98.3%
98.2%
96.0%
94.1%
91.2%

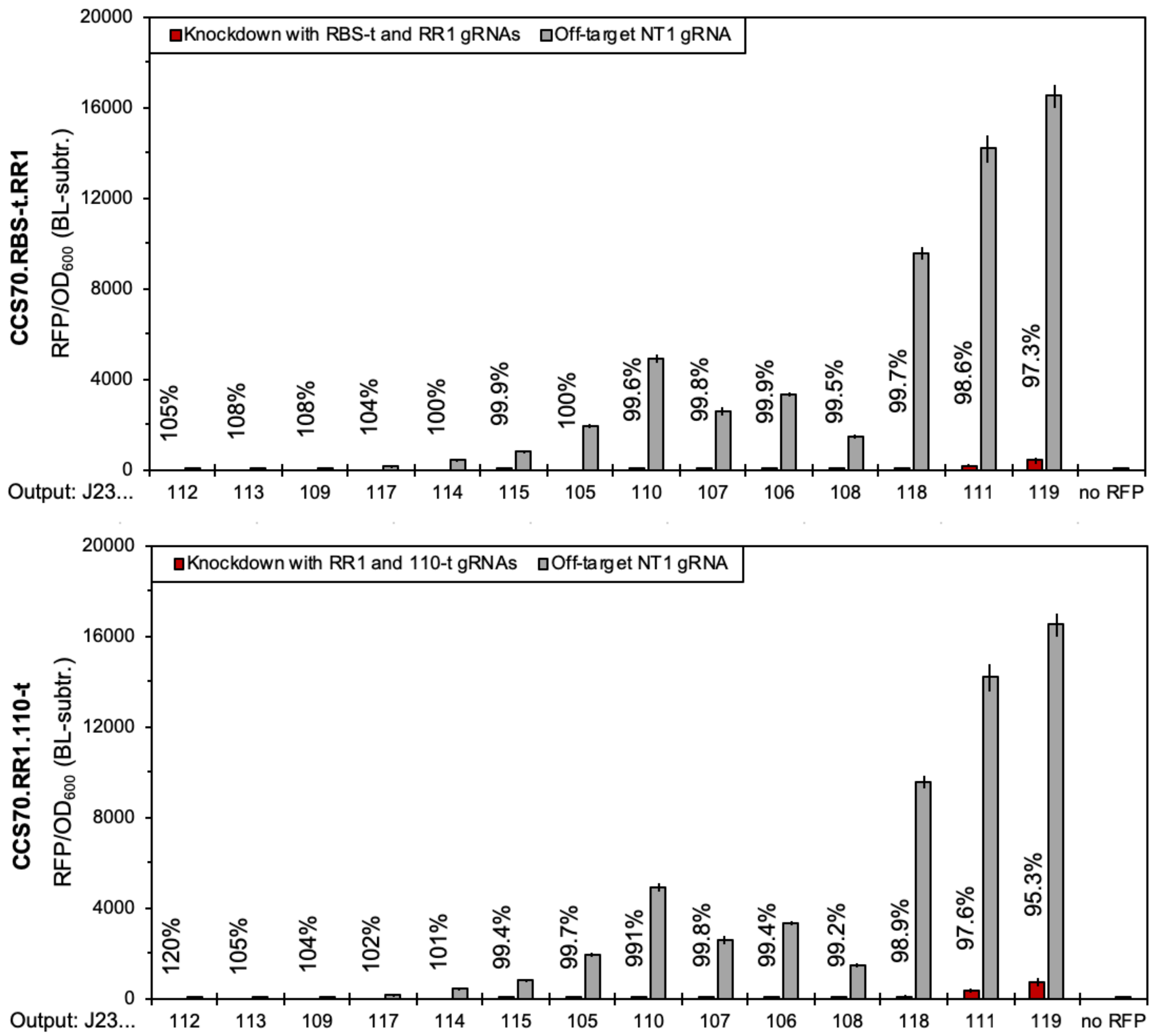


Supplementary Figure 3. Knockdown levels of Anderson promoters in additional CCS strains not shown in Figure 3 in the main text. Grey bars indicate RFP expression from each promoter in the absence of knockdown due to an off-target gRNA, and red bars indicate repressed expression due to the gRNA(s) indicated in the strain name on the y-axis. Percentages indicate knockdown levels.

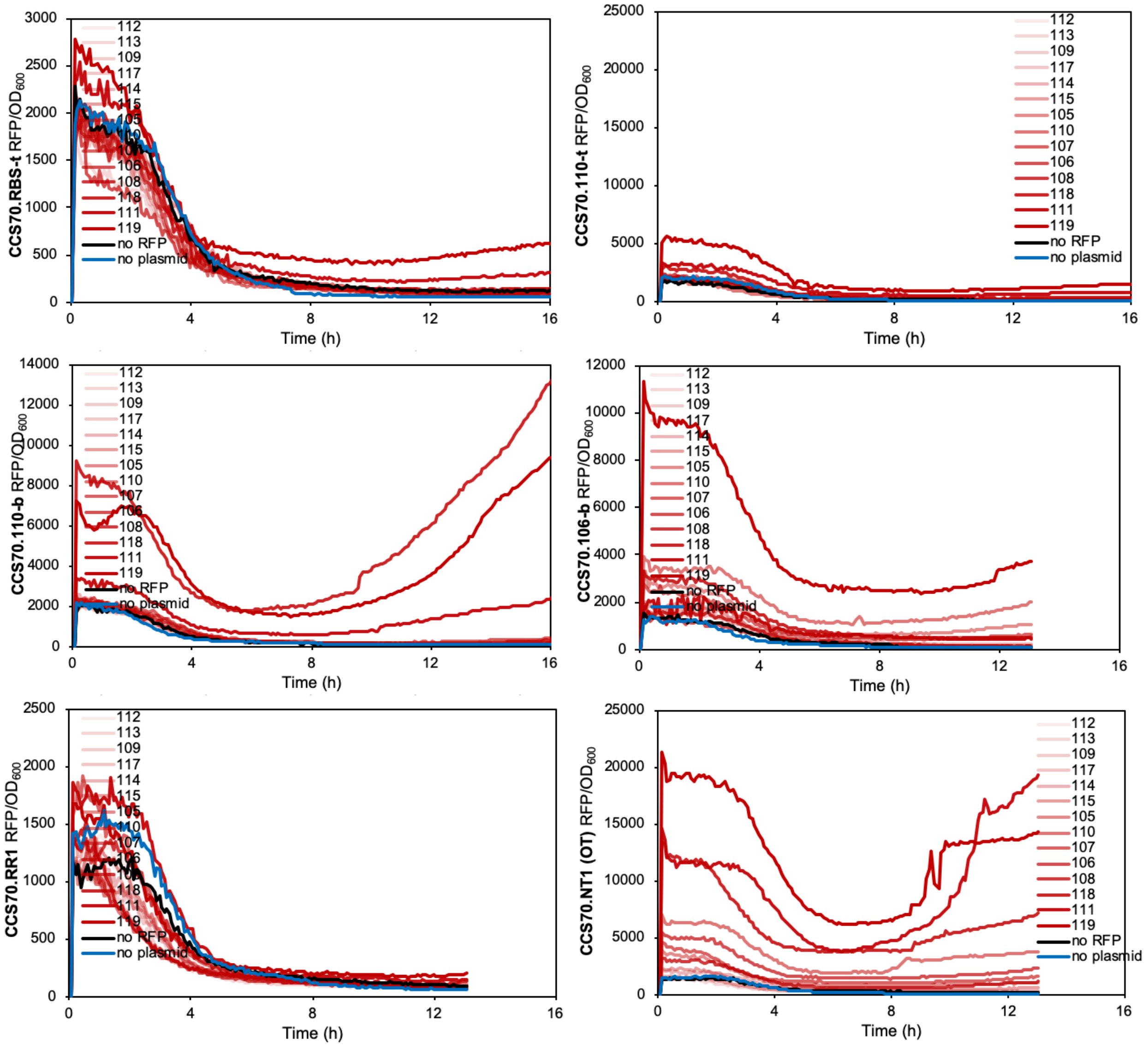


Supplementary Figure 4. Kinetic RFP expression data from the Anderson promoter series in each of the CCS strains that target a subset of that series.

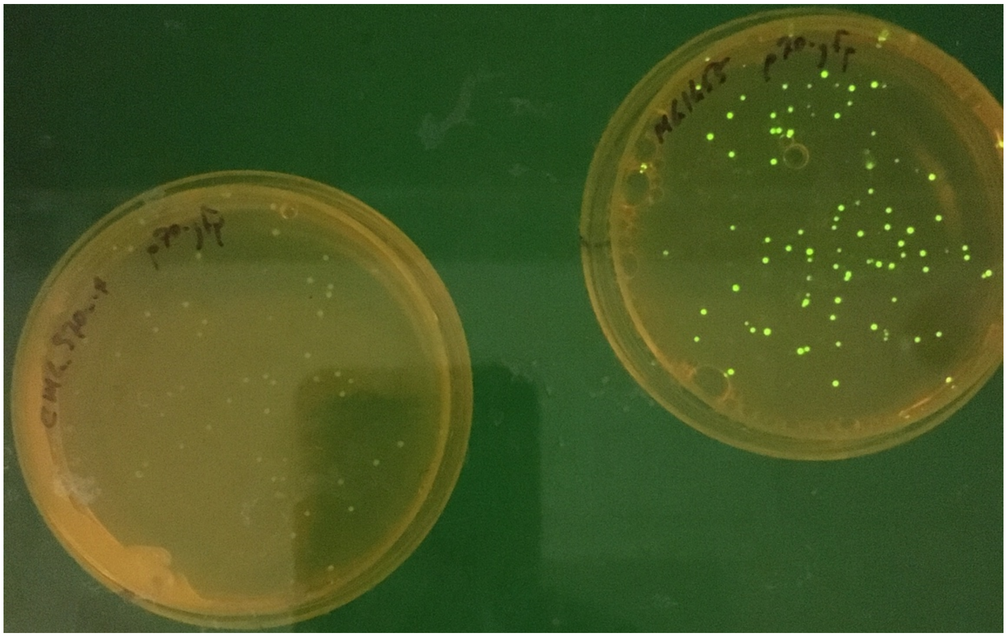

Supplementary Figure 5.  Knockdown of the σ70 promoter by the S70a-t gRNA (left) only in colonies on agar plates (right plate, no CRISPRi).

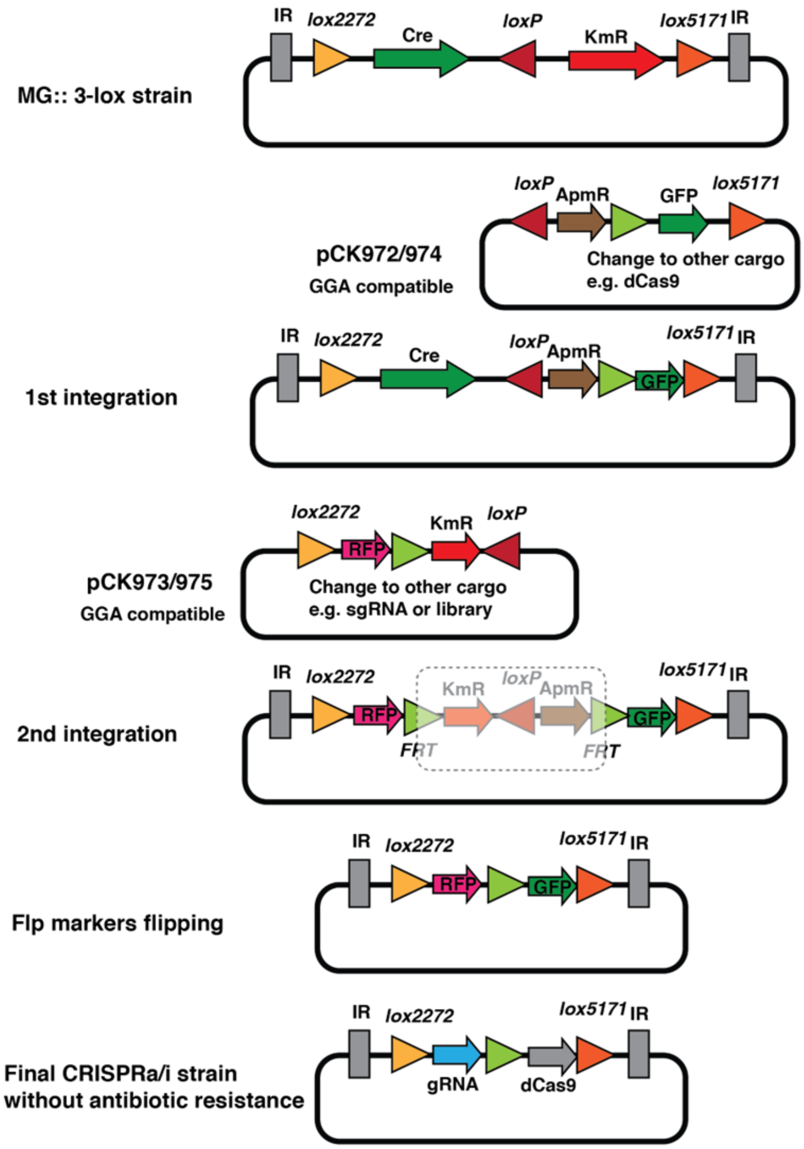


Supplementary Figure 6. General curable CRAGE-Duet workflow, here delivering GFP with apramycin resistance as the first cargo and RFP with kanamycin resistance as the second cargo, then removing both resistance markers with FLP recombinase.
